# A small cationic probe for accurate, punctate discovery of RNA tertiary structure

**DOI:** 10.1101/2025.08.29.672938

**Authors:** Jeffrey E. Ehrhardt, David Y. Qiu, Shouhong Jin, Mark A. Boerneke, Caroline J. Aufgebauer, Stacy M. Horner, Kevin M. Weeks

**Author notes:** contributed equally (Jeffrey E. Ehrhardt, David Y. Qiu).

## Abstract

RNA molecules fold into intricate three-dimensional tertiary structures that are central to their biological functions. Yet reliably discovering new motifs that form true tertiary interactions remains a major challenge. Here we show that RNA tertiary folding occasionally generates electronegative motifs that react selectively with the small, positively-charged probe trimethyloxonium (TMO). Sites with enhanced reactivity to TMO, compared with the neutral reagent dimethyl sulfate (DMS), are indicative of tertiary structure and define T-sites. These positions share a structural signature in which a reactive nucleobase is adjacent to non-bridging phosphate oxygens, creating localized regions of negative charge. T-sites consistently map to the cores of higher-order structural interactions and functional centers across diverse RNAs, including distinct states in conformational ensembles. In the 10,723-nt dengue virus genome, three strong T-sites were detected, each within a complex structure required for viral replication. Cation-based covalent chemistry enables high-confidence discovery and analysis of functional RNA tertiary motifs across long and complex RNAs, opening new opportunities for transcriptome-wide structural analysis.

## Introduction

RNA molecules encode information through both primary genetic sequence and via higher-order structural elements. RNA elements with complex structures mediate biological functions ranging from catalysis, to ligand responses, to translation regulation^1–4^. Chemical probing has been used extensively to characterize and define RNA structures, and recent developments in massively parallel sequencing have enabled high-throughput characterization of RNA secondary structure. In particular, methods using SHAPE and DMS probing with read out based on mutational profiling (MaP) and single-molecule correlated chemical probing (smCCP) strategies are revealing that coding and non-coding RNAs contain extensive, often directly functional, internal secondary structure elements that are expressed both as stable structures and as ensembles^5–7^. Experimentally-informed secondary structure modeling, including for large and complex RNAs, is now well established and typically highly accurate^6,8–10^. Chemical probing is now an essential tool for understanding RNA structure-function interrelationships at the level of RNA base pairing, extending from understanding the regulatory logic of RNA viruses, to defining how RNA structure functions as a ubiquitous switch or rheostat in bacterial and eukaryotic systems, to optimizing RNA medicines^5,6,11–13^.

By contrast, the extent to which true *tertiary* structures form in RNAs remains poorly explored. RNA tertiary structures are highly diverse and include non-canonical pairing, kissing loop contacts, pseudoknots, and indirect backbone interactions^14,15^. The structural diversity of tertiary interactions and the fact that even complex tertiary structure motifs contain extensive simple base pairing has made devising broadly applicable chemistry for definitive RNA tertiary structure discovery challenging. Conventional chemical probing technologies are certainly sensitive to formation of tertiary contacts, as these methods generally measure conformational flexibility or steric accessibility. However, these approaches do not distinguish whether a given reactivity measurement or difference is reflective of base pairing versus higher-order tertiary interactions. Existing probing strategies that show partial selectivity for RNA tertiary structures have done so via measurements of solvent accessibility, crosslinking (UV, psoralen or SHAPE based), reaction with soft metal ions (for example, Pb^2+^ and Tb^3+^), or single-molecule correlated chemical probing^7,16–18^. Drawbacks of extant approaches include low signal above background, high read depth requirements when read out by massively parallel sequencing, and convolution of secondary and tertiary structure features. Current strategies are thus limited due to challenging implementation, difficulty assigning measurements specifically to RNA tertiary structure, and lack of suitability for *de* novo RNA tertiary structure discovery.

Here we devise a chemistry-based probing strategy, highly selective for RNA tertiary structure, by taking advantage of structure and charge features unique to RNA motifs with complex folds. RNAs with complex structures create surfaces with distinctive electrostatic profiles, which result from confining the negative charge in the phosphoribose backbone in tightly folded tertiary motifs^15,19,20^. We developed a comparative chemical probing strategy that measures reactivity differences between positively-charged and neutral reagents. In probing an ∼11,000-nt dengue virus (DENV) RNA genome, we identified three strong tertiary structure elements: two show the expected electronegative signature based on comparison with known structures, and the third corresponds to a proven regulator of viral replication. This cation-based chemical probing strategy thus identifies distinctive electronegative motifs in a precise and punctate way, enabling detection of RNA tertiary structures with high accuracy and low false-positive discovery.

## Results

### Differential reactivity of TMO and DMS for RNAs with complex structures

We reasoned that the key to developing a chemical probing strategy highly selective for true RNA *tertiary* structure would be to focus on RNA electrostatics. Individual RNAs feature diverse backbone geometries, extending from the smooth helix characteristic of simple helices to tight kinks and turns found in riboswitches and RNAs with complex structures^15^. Specific unusual conformations of an RNA backbone might create an electrostatic environment favoring or disfavoring reaction by charged molecules.

We recently introduced trimethyloxonium (TMO) as a reagent for RNA structure probing^21^. TMO is similar in size to and alkylates RNA via the same electrophilic mechanism as dimethyl sulfate (DMS) (Figure 1). However, TMO has a shorter half-life than DMS and is positively charged. The short half-life of TMO can be advantageously leveraged to study time-resolved RNA folding and to capture through-space interaction networks during folding^21^. Here we will leverage the positive charge on TMO to create a chemistry-based strategy for RNA tertiary structure discovery. TMO and DMS reactivities are efficiently quantified using massively parallel sequencing via mutational profiling (MaP) reverse transcription^21,22^, including at low read depths (Figure S1).

**Figure 1.**
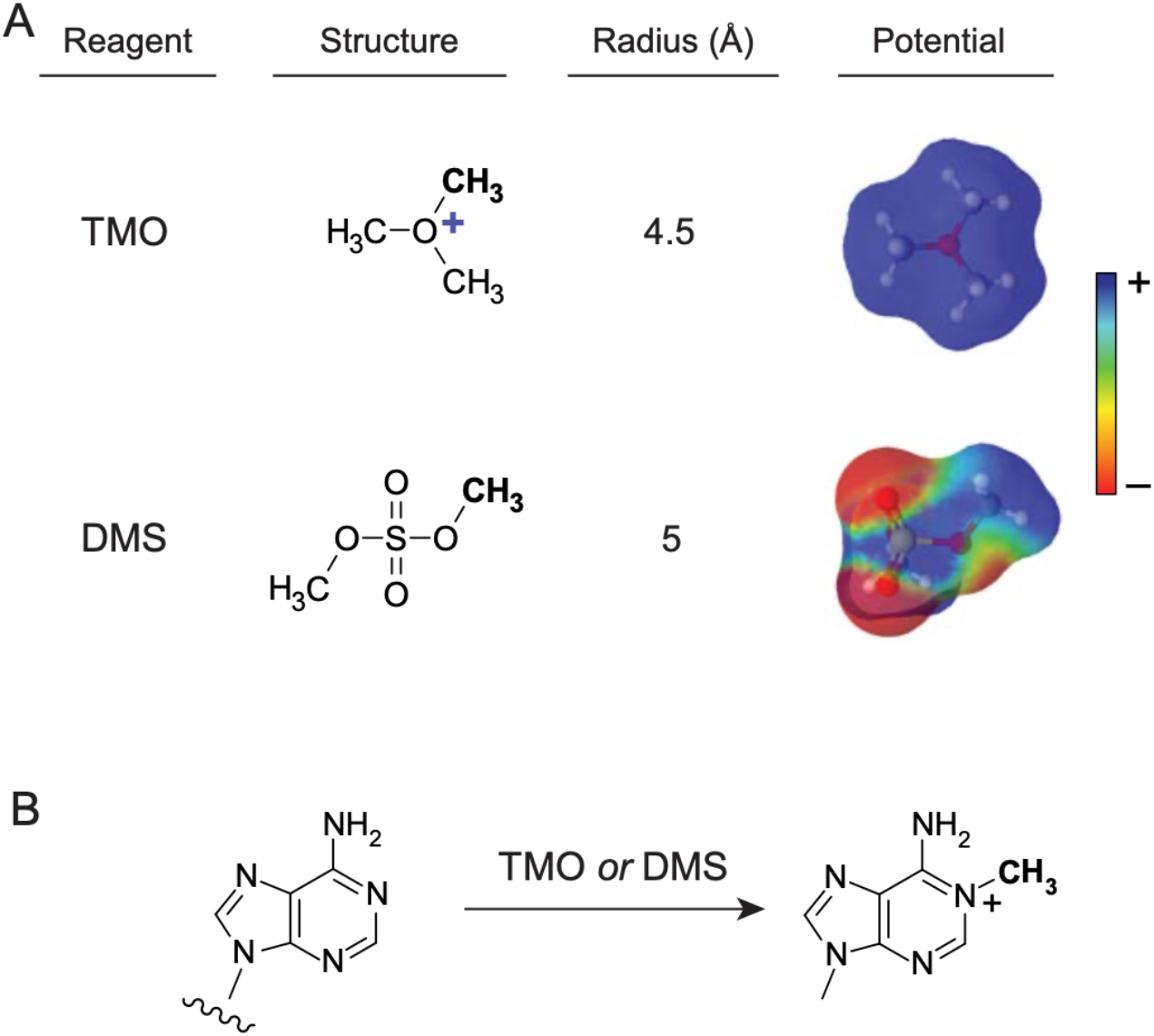
Characteristics and RNA chemistry for TMO and DMS. (**A**) Structures and surface charge. (**B**) Reaction with adenine.

We probed three RNAs of known three-dimensional structure: the catalytic core of a bacterial RNase P (termed RNase P), the thiamine pyrophosphate (TPP) aptamer domain from the TPP riboswitch, and a group II intron. All three RNAs contain complex tertiary structures critical to their biological function^23–25^. RNase P cleaves tRNAs during their biogenesis, the TPP riboswitch binds TPP to regulate gene expression in prokaryotes, and group II introns are self-splicing ribozymes. Across all three RNAs, the reactivity patterns of TMO and DMS reactivity were broadly similar. Both probes reacted with nucleotides in unstructured loop and bulge regions, consistent with their expected reaction mechanism with RNA (Figures 1B, 2A).

Strikingly, we observed distinctive, large-magnitude, TMO-specific reactivity at a single site in each model RNA. In each case, the differentially reactive nucleotide was an adenosine. In RNase P, the largest difference in reactivity was at adenosine 45, where TMO was ∼20-fold more reactive than DMS (Figure 2C). Although nucleotide 45 is unpaired in secondary structure diagrams, it is structurally constrained in the catalytic core of the RNase P RNA by base stacking and non-canonical interactions (Figure 2A)^24^. In the TPP aptamer domain at nucleotide A44 and in the group II intron at nucleotide A376, we observed 10- and 8-fold increases in reactivity of TMO over DMS, respectively (Figure 2B, 2D). As with RNase P, these sites are in the complexly folded tertiary core of each RNA^23,25^. We refer to these <u>T</u>MO-reactive sites, indicative of RNA tertiary structure, as T-sites.

**Figure 2.**
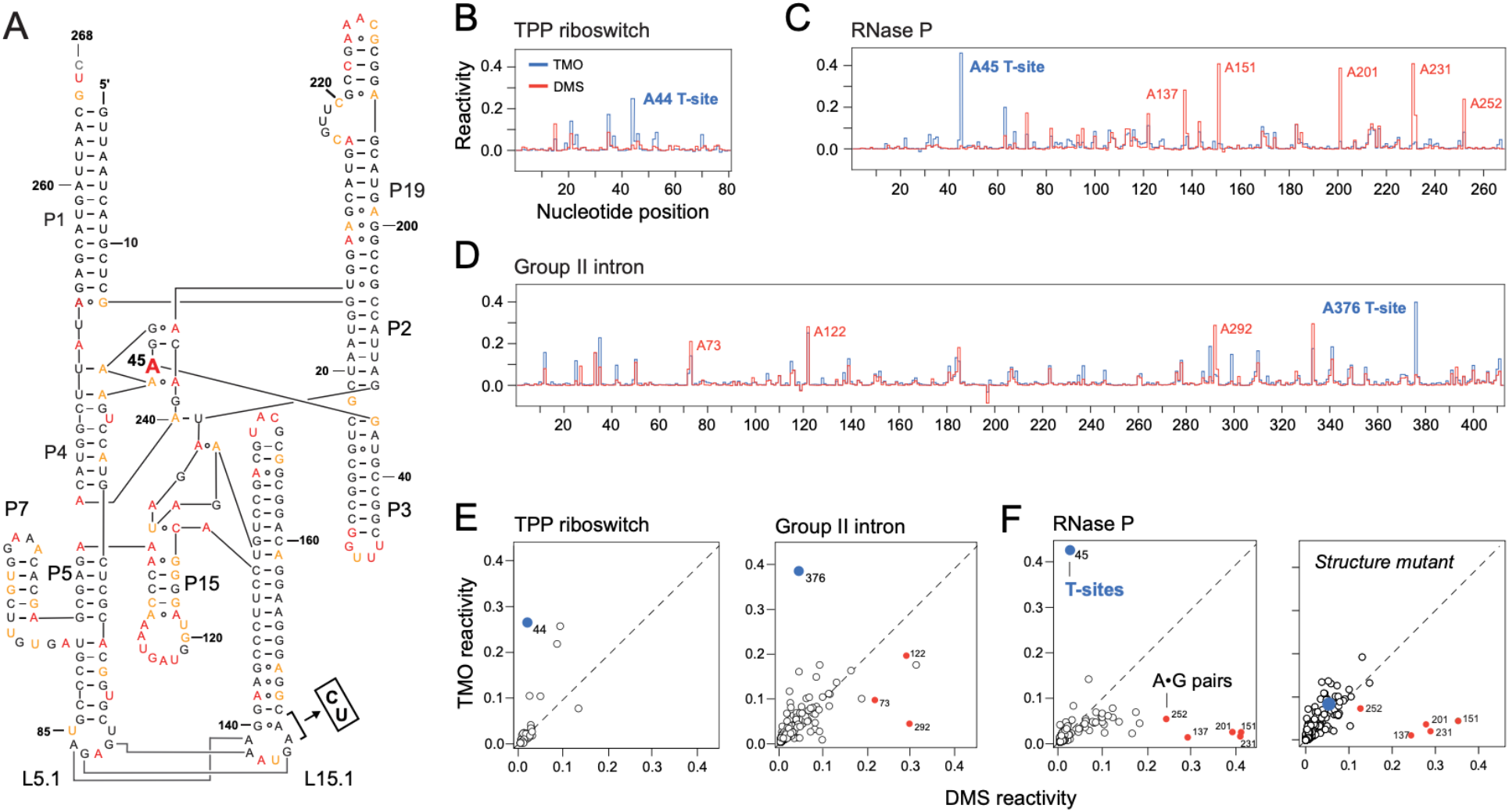
TMO and DMS reactivity profiles for three RNAs. (**A**) Secondary structure of RNase P. T-site (at position 45) is shown as large bold A. Nucleotides are colored by TMO reactivity; red, orange and black correspond to high, medium and low reactivities, respectively. Secondary structure is annotated with a mutation (AA to CT in L15.1, *box*) that selectively disrupts tertiary structure^21^. (**B-D**) TMO (blue) and DMS (red) reactivity profiles for (**B**) TPP aptamer domain, (**C**) RNase P, and (**D**) group II intron. Sites with significant differential reactivities are labeled; T-sites are in bold blue text; sites with DMS reactivity greater than TMO reactivity (A•G pairs) are labeled in red. (**E**) Correlation plots of reactivity rates for RNA modified with DMS (x-axis) and TMO (y-axis). T-sites (blue) and A•G pairs (red) with distinctive reactivities are labeled. (**F**) For RNase P, both the native sequence and the tertiary structure-disrupting mutant are shown.

We also observed the converse: At a subset of adenosine residues in the RNase P and group II intron RNAs, DMS was more reactive than TMO (Figure 2C, 2D). We will show below that these DMS-reactive sites correspond to adenosines involved in A•G base pairs. These differential TMO and DMS reactivity signals are distinctive in that one reagent generally shows low reactivity, while the other is highly reactive (Figure 2E, 2F). These patterns imply that there is a structure-based mechanism that creates specific probe-binding interaction sites that govern each class of reactivity.

### T-site reactivities are strongly and specifically dependent on RNA tertiary structure

There is a single T-site in each of the model RNAs, which span a wide range in size (80 to 412 nts) and have completely different global architectures. The reactivity signal was separated from background for each RNA by large factors, among or exceeding the largest we have ever observed in chemical probing experiments (Figure 2E, 2F). These features suggest that T-sites are strong predictors of specific RNA tertiary structure motifs.

We initially examined the hypothesis that T-sites are strictly dependent on formation of higher order RNA structure in two ways. First, we mutated two nucleotides in the L15.1 loop in RNase P to weaken a loop-loop interaction that is critical to higher-order folding of this RNA^21,24^ (Figure 2A). As expected, nucleotide reactivities increased precisely at the (L5.1-L15.1) loop-loop interaction, consistent with its disruption (Figure S2A). Probing with TMO and DMS showed that the overall reactivity pattern for each reagent was otherwise largely unperturbed, consistent with maintenance of the native complex base-paired secondary structure. By contrast, differential TMO-vs-DMS reactivity was lost at the T-site, reflective of selective loss of tertiary structure in this mutant (Figure 2F). In sum, T-site reactivity, but not general per-nucleotide reactivity, is specifically and strongly dependent on formation of native tertiary structure in RNase P.

Second, we examined TMO reactivities for the TPP aptamer domain, RNase P and group II intron RNAs as a function of Mg^2+^, which stabilizes closely packed negatively charged helices in RNA tertiary structures. All three RNAs adopted native-like RNA secondary structure folds in the absence of Mg^2+^, as supported by the correlation between per-nucleotide reactivities in the presence and absence of Mg^2+^. In contrast, the T-site signal, due to TMO, is substantially reduced when the three RNAs were probed in the absence of Mg^2+^ (Figure S2B). These observations are consistent with the model that a specific, constrained higher-order tertiary structure enables an experimentally observable T-site.

### T-site and A•G base pair structure determinants

Each of the T-sites we observed occurs in a region of closely packed RNA strands (Figure 3A). The T-sites in the three RNAs are structurally distinct, but each site shares specific atomistic features (Figure 3B). First, the reactive T-site in each RNA is an unpaired adenosine. Second, local folding of the RNA backbone places the structurally constrained adenosine residue within 4-7 Å of two nonbridging phosphate oxygen atoms. Third, the reactive adenosine residue of each T-site stacks with nearby nucleobases. Despite these atomistic similarities, the specific arrangements of the RNA strands in the three RNAs are completely different (Figure 3B). Indeed, in the three RNAs studied here, the sequence distance between the T-site adenosine and a non-bridging oxygen varies from 30 nucleotides in the TPP aptamer domain to 371 nucleotides in the group II intron. There are thus a wide variety of local RNA geometries capable of creating a T-site, but such sites occur only in regions of complex tertiary interactions. These features support a chemical model in which TMO preferentially binds at an electronegative interaction site adjacent to the reactive adenosine N1 atom (Figure 3C).

**Figure 3.**
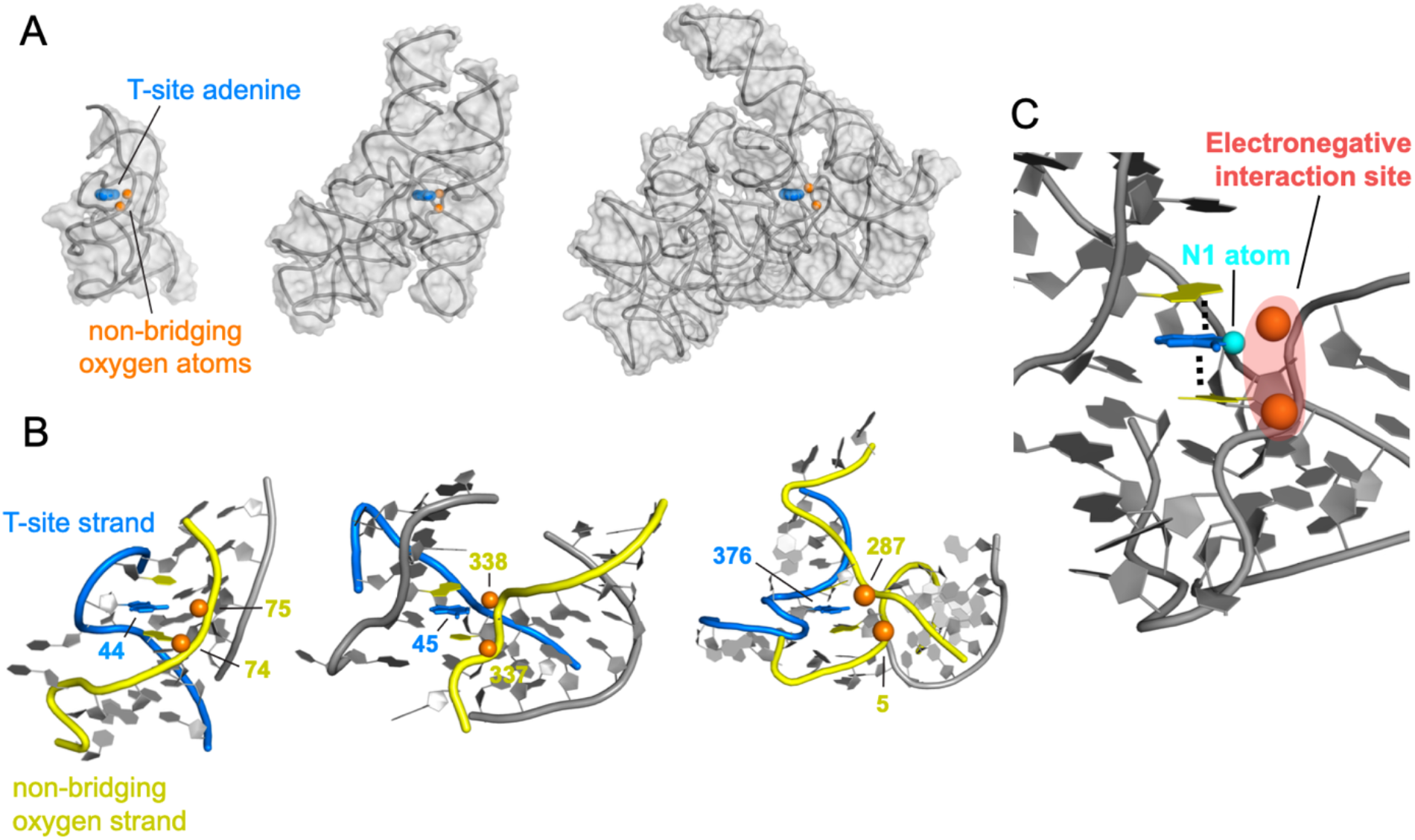
Visualization of T-sites in RNAs with complex tertiary folds. (**A**) RNA structures (grey) showing T-site residue (blue) situated proximal to nonbridging phosphate oxygen atoms (orange spheres). (*left* to *right*) TPP riboswitch aptamer domain, RNase P and group II intron. RNAs are positioned such that T-site adenosine residues have the same orientation. (**B**) Detailed views showing geometry of the T-site adenine nucleobase (blue), proximal nonbridging oxygen atoms (orange), and nucleotides that stack with the reactive position (yellow). Sequence positions of T-site adenosine and proximal non-bridging oxygen atoms are numbered. RNA backbones for the strands on which the T-site adenosine and non-bridging phosphate atoms are located are emphasized in blue and yellow, respectively. (**C**) Summary of features found at T-sites. T-site adenosine is blue; reactive N1 atom is cyan; the two closest non-bridging oxygens are orange; and nucleotides that engage in base stacking with the T-site adenine nucleobase are yellow. Region of electronegative potential is emphasized in red.

In addition to the single adenosine residue in each RNA that was preferentially reactive with TMO, we also identified multiple sites in the RNase P and group II intron RNAs that showed the converse behavior and reacted more strongly with DMS than with TMO (Figure 2). These sites all occur at A•G base pairs^24,25^ (Figure S3), one of the most common non-canonical pairs^26,27^. All pairs that are preferentially reactive toward DMS are *trans* A•G pairs that interact via the Hoogsteen face of A and the sugar edge of G. This pairing geometry creates an adenosine nucleotide with a distinctive conformation in which the adenosine N1 atom is accessible for reaction and is juxtaposed near the 2’-OH of the pairing G residue (Figure S3). We hypothesize that this 2’-OH group facilitates optimal positioning of DMS, which is predictive for A•G non-canonical base pairing and allows their identification in complex RNAs.

### Mapping higher-order RNA structure around T-sites

To understand the relationship between T-sites and higher-order structure, we assessed the effect of disrupting, by mutation, RNA tertiary structure specifically at each T-site. For each model RNA, we mutated the T-site adenosine to uridine or introduced 1- or 3-nucleotide deletions centered at the T-site. The overall reactivity profiles for each of the mutants were similar to those of the native sequence when the RNAs were probed with TMO, emphasizing that the mutants adopt native-like RNA secondary structure. However, specific nucleotides within each RNA mutant displayed punctate increases in reactivity (Figure 4A and S4). The nucleotides with increased reactivities were not involved in canonical base pairing interactions and all lie close in three-dimensional space to the mutated T-site but were often distant primary sequence (Figure 4B). In sum, analysis of RNA T-site disruption mutants reveals the close-in-space environment surrounding the T-site and further supports the model that T-sites report critical, core interactions in an RNA tertiary structure.

**Figure 4.**
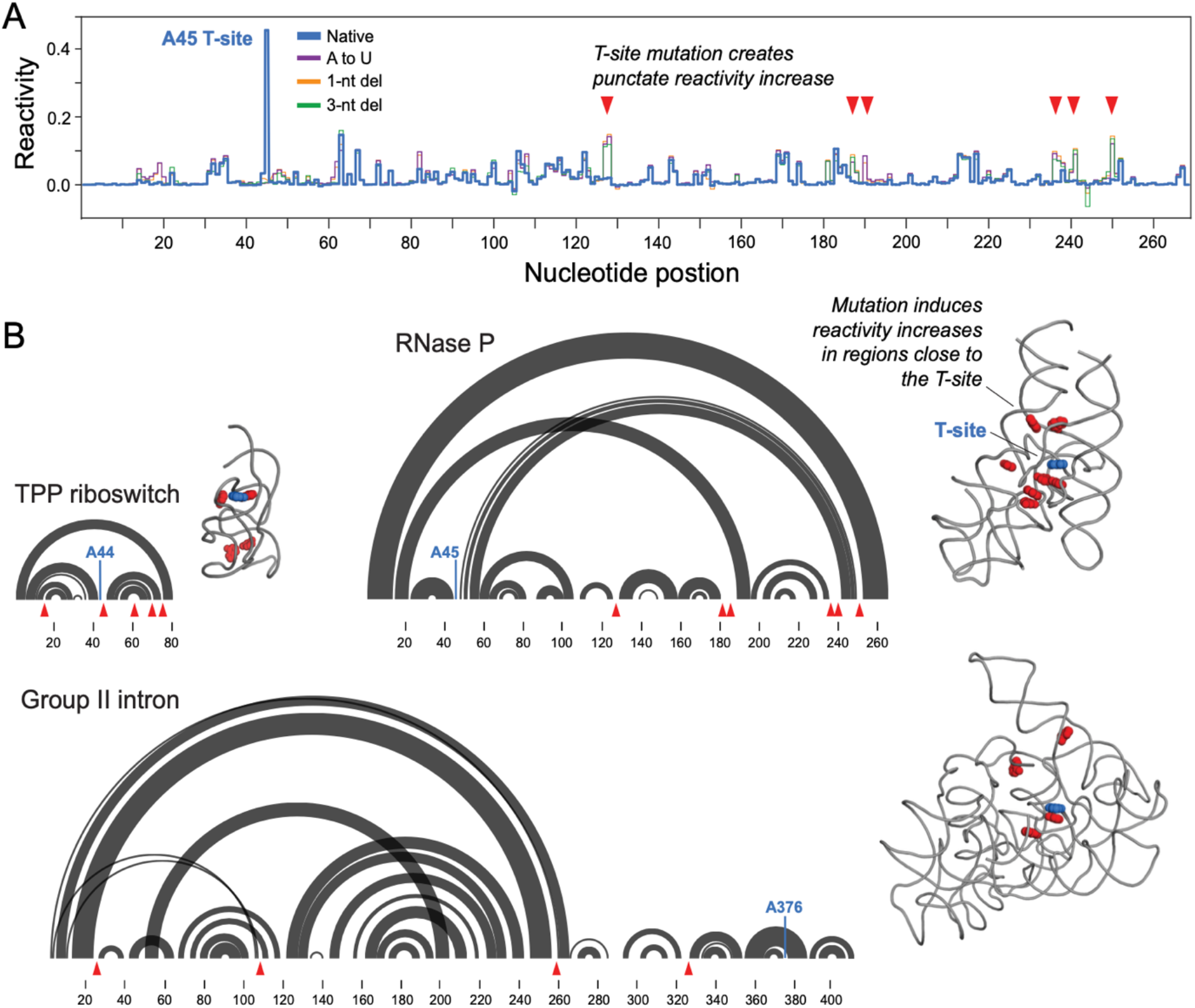
Mapping the structural neighborhood surrounding a T-site. (**A**) Comparison of TMO reactivity profiles for native and mutant sequences for the RNase P RNA. Nucleotides that show significant (≥0.05) increases in reactivity rate in the 3-nt deletion mutation are denoted with red triangles. (**B**) Secondary and tertiary structures for the TPP aptamer domain, RNase P, and group II intron RNAs, the locations of T-sites, and perturbations induced by mutating the T-site. Secondary structures (base pairing) are shown as arcs (grey). Three dimensional structures shown as gray backbone. T-sites are labeled in blue. Nucleotides that show significant reactivity increases in 3-nt deletion mutants are emphasized in red. Reactivity profiles for the TPP riboswitch and group II intron are shown in Figure S4.

### T-sites identify the tertiary structure-containing state in a conformational ensemble

Many RNAs populate conformational ensembles, as exemplified by riboswitches^28^. Riboswitches typically contain a ligand-sensitive aptamer domain and an adjoining expression platform (Figure 5A). The ThiM riboswitch populates translational OFF and ON states, where the OFF state is stabilized when the TPP ligand binds the aptamer domain, which has the well-defined tertiary structure characterized above (Figure 3A). The OFF state also sequesters the Shine-Dalgarno (SD) site, essential for translation initiation, in a stable helix^29^. As demonstrated here, this state contains a T-site at nucleotide 44 (Figure 5A). The ON state adopts a fold containing simple stem-loop motifs and that presents the start codon and Shine-Dalgarno sequence in a single stranded region, accessible for interaction with the ribosome and initiator tRNA (Figure 5A). The OFF state is populated in both the presence and absence of TPP, at ∼80 and 40%, respectively, for our structure-switching riboswitch construct (Figure S5).

**Figure 5.**
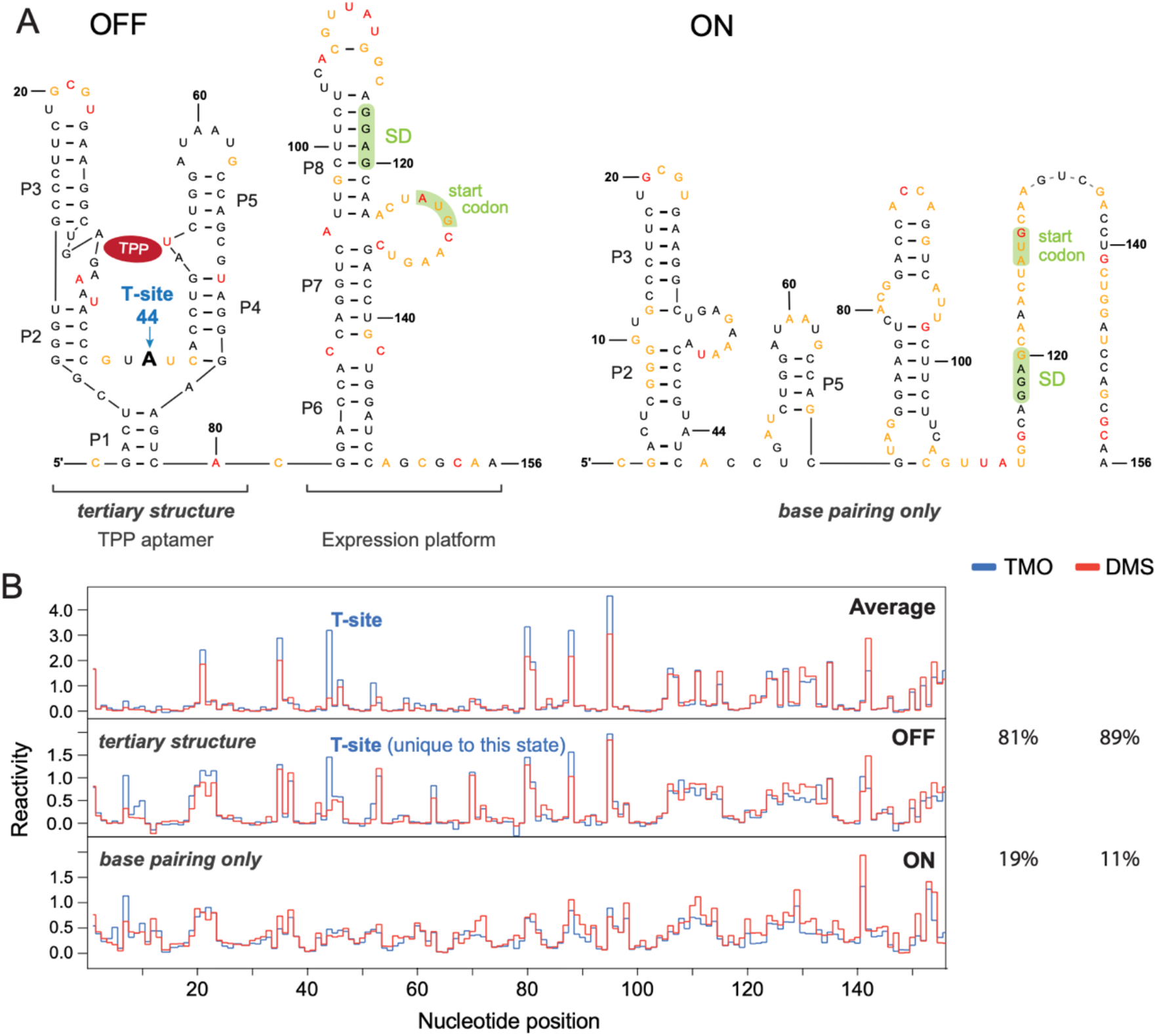
Detecting tertiary structure specific to individual state in a conformational ensemble for a structure-switching TPP riboswitch. Riboswitch was probed in the presence of saturating concentrations of the TPP ligand. (**A**) Secondary structures of the ThiM TPP riboswitch in the OFF and ON translation states, modeled by DANCE-MaP based on DMS-MaP data. Nucleotides are colored by DMS reactivity. Red, orange and black correspond to high, medium, and low reactivities, respectively. T-site (at position 44) is shown as large bold A; the Shine–Dalgarno (SD) and translational start codon are highlighted with green. (**B**) TMO (blue) and DMS (red) reactivity profiles for the TPP riboswitch. The upper panel shows the averaged, bulk reactivities, prior to deconvolution. Middle and lower panels show reactivities deconvoluted by DANCE-MaP^31^. Population percentages for the OFF and ON states, determined from deconvolution of TMO and DMS data, respectively, are listed at right. Comparative data, obtained in the presence and absence of TPP, are provided in Figure S5.

We probed the structure-switching riboswitch RNA in the presence and absence of TPP with both TMO and DMS. We used a single-molecule correlated chemical probing (smCCP) strategy, called DANCE-MaP, to deconvolute multiple conformations in the riboswitch ensemble^30,31^. As expected, bulk RNA reactivities showed a T-site at position A44 in the presence of TPP (Figure 5B, *top*). Ensemble deconvolution, based on either reagent, yielded similar populations of OFF and ON states (Figure 5B, *right*). Based on the secondary structures modeled for each state by DANCE, we assigned the SD-sequestered configuration as the OFF state. We could then recreate the reactivity patterns for each state, for each reagent. We observed the T-site only in the OFF state, which therefore correctly identifies the state with the TPP-binding *tertiary* structure (Figure 5B). Strikingly, in the absence of TPP, where the OFF state comprises only ∼40% of the population, the T-site is not visible in the bulk per-nucleotide reactivity profiles but becomes identifiable in the deconvoluted OFF state (Figure S5). In sum, T-site probing specifically identifies the state in a conformational ensemble that contains higher-order tertiary structure and can reveal an, otherwise hidden (low population), tertiary structure in a complex ensemble.

### Tertiary structure discovery across a viral RNA genome

Viral RNA genomes contain a diverse array of complex structures, and indeed encompass a large fraction of all known RNA tertiary structure motifs relevant to eukaryotic organisms^4^. We probed DENV serotype 2 (DENV2, New Guinea C strain) RNA, extracted from virions, with TMO and DMS. We obtained high-quality data across the entire 10,723-nt genome and identified three strong T-sites at positions A8293, A10417, and A10473 (Figure 6A; and Figure S6 for the entire genome). Strikingly, all three sites play pivotal roles in DENV2 replication. Two sites lie in the 3′-UTR, a region densely populated with replication control elements and RNA tertiary structures^4,32^. These two sites, A10417 and A10473, reside within an exonuclease resistant RNA (xrRNA) motif and a Dumbbell motif, respectively.

**Figure 6.**
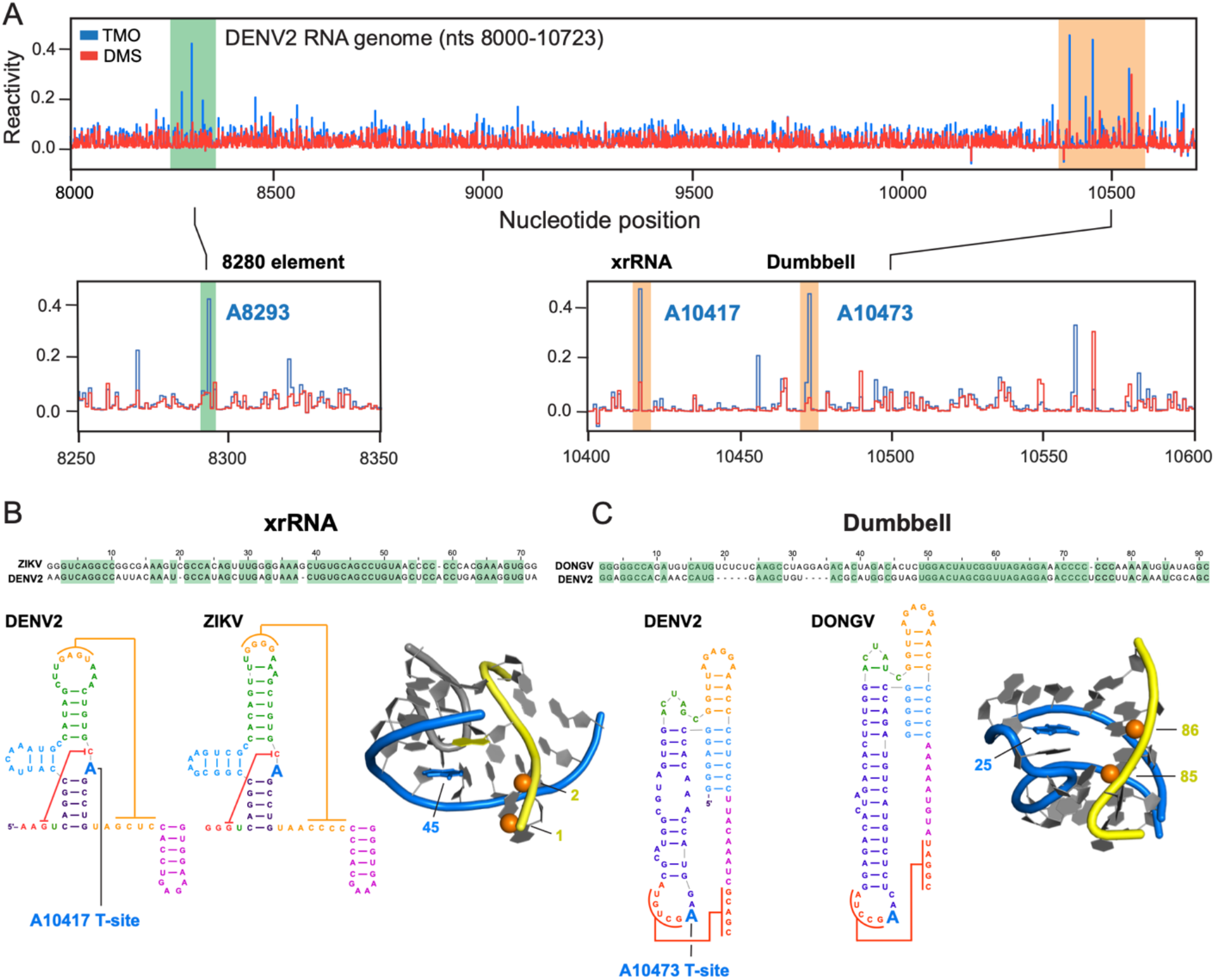
T-site detection across the DENV2 RNA genome. (**A**) Reactivity profiles for TMO (blue) and DMS (red) over a representative region of the DENV2 genome. Regions containing T-sites are shaded, and T-sites are labeled. Sequence alignments, secondary structure models, and three-dimensional models for (**B**) xrRNA and (**C**) Dumbbell motifs. DENV2 structures are compared with their ZIKV and DONGV analogs, for xrRNA and Dumbbell motifs, respectively. T-site nucleotides are emphasized with large blue nucleotide lettering. For the three-dimensional models, the T-site nucleotide and non-bridging phosphate oxygen atoms are shown in blue and as orange spheres, respectively; RNA backbones for the strands on which the T-site adenosine and non-bridging phosphate atoms are located are emphasized in blue and yellow, respectively. Nucleotides are numbered based on the crystal structures^33,34^. Full TMO and DMS probing data across the DENV2 genome are shown in Figure S6.

xrRNA and Dumbbell motifs are found across multiple viral superfamilies and representative structures of these elements have been determined^33,34^. xrRNA motifs form a highly stable, pseudoknot-containing structure, which prevents degradation by cellular 5’⟶3’ exonucleases^35^. The DENV2 xrRNA is critical for viral replication^36^ and its sequence is highly conserved with that of the Zika virus (ZIKV), whose structure is known (Figure 6B). Examination of the adenosine nucleotide in ZIKV^33^ corresponding to the DENV A10417 T-site shows that this nucleotide, indeed, adopts the precise T-site conformation. The adenosine N1 atom points toward non-bridging phosphate oxygen atoms from a nearby backbone (Figure 6B). Dumbbell motifs, including that in DENV^37^, regulate the transition from viral replication to translation. Analysis of a homologous Dumbbell structure from a related insect-specific flavivirus, Donggang virus (DONGV)^34^, corresponding to the DENV A10473 T-site, again, reveals this nucleotide forms a T-site geometry (Figure 6C). For the third T-site, at position A8293, no 3D structure is known. However, the motif, termed the 8280 element, is conserved among all four DENV serotypes and mutations near A8293 significantly decrease viral replication and infectivity, highlighting the biological significance of this T-site motif^38^. These results emphasize that T-site probing enables *de novo* detection of RNA tertiary structure, across long RNAs, in a highly specific and punctate manner.

Both the xrRNA and Dumbbell motifs occur twice in the DENV2 3′-UTR, and we detect only one T-site instance for each element. The first xrRNA motif (we detect the second) has a uridine instead of an adenosine nucleotide at the T-site position, and T-site chemistry (Figure 1B) currently only detects tertiary structure at adenosine residues. The potential T-site (position A10561) in the second Dumbbell motif (we detect the first) does display moderate TMO over DMS reactivity (Figure 6A), but not large enough in magnitude to meet stringent T-site thresholds defined from model 3D RNA structures (see Methods). We posit that the tertiary structure in the second Dumbbell is less stable than in the first, resulting in the lower differential reactivity. The presence of a T-site is thus highly predictive for RNA tertiary structure, but the absence of a T-site does not imply absence of higher-order structure.

## Discussion

The key to precisely mapping RNA *tertiary* structure using chemical tools lies in finding a measurement that reflects a through-space interaction strongly correlated with higher-order structure. Absent such an insight, efficient *de novo* tertiary structure discovery has remained an important, widely sought, but unrealized goal. Here we show that differences in per-nucleotide reactivities for TMO and DMS, similar-in-size reagents with different charges, identify electronegative motifs present in specific RNA tertiary structures. Our chemical probing strategy is unique in its ability to measure long-range, through-space interactions – between an adenosine N1 atom and distantly positioned non-bridging oxygen groups – via an individual-nucleotide reactivity measurement (Figure 3C).

Multiple lines of evidence emphasize that T-site chemistry is highly selective for RNA motifs that form true tertiary structure motifs. First, the T-site in RNase P disappears for a point mutant^21^ that disrupts the tertiary structure but does not change the complex secondary structure (Figure 2F and S2A). Second, T-sites in the TPP riboswitch, RNase P and group II intron RNAs are disfavored when TMO and DMS probing are performed in the absence of Mg^2+^ ion (Figure S2B). Third, mutation of a T-site induces close-in-space changes in TMO reactivity, consistent with disruption of local tertiary structure (Figure 4 and S4). Fourth, for a structure-switching TPP riboswitch construct, which forms a two-state ensemble in which one state forms a tertiary structure and the other state just contains base pairing^31^, we only observe a T-site in the OFF state that forms true higher-order structure (Figure 5B and S5).

We identified a single T-site in each of three highly structured RNAs with known tertiary structures, plus three strong T-sites across the DENV RNA genome. We also observed eight sites with the reverse behavior – enhanced DMS reactivity – predictive of sheared A•G Hoogsteen-ribose face base pairs, one of the most common non-canonical pairing interactions^26,27^. The dual probing experiment chemistry is simple and direct, and read-out employs the efficient, well-validated MaP strategy^21,22,39^. Data interpretation is straightforward; the raw difference in reactivity rates between TMO and DMS yields the T-site signal. This differential signal is among the largest we have observed in a chemical probing experiment and is readily detectable at low sequencing read depths, opening the door for applications on transcriptome-wide scales. By introducing a mutation at a T-site (we favor the 3-nt deletion), the through-space structural neighborhood surrounding the T-site is defined as punctate increases in reactivity due to loss of tertiary interactions. The strong signal-to-background and rarity of T-site signals indicates this strategy can be broadly applied for *de novo* RNA tertiary structure discovery.

T-sites are created by motifs that form unusual, but stable, local electronegative interaction sites immediately adjacent to an accessible adenosine N1 atom (Figure 3C). This precise geometry is formed from highly diverse tertiary structure folds. Indeed, in the five T-sites visualized here (Figures 3 and 6), there is no similarity in backbone geometry or RNA strand orientation, beyond the proximal positions of the reactive adenine nucleobase and non-bridging oxygen atoms. In RNase P and the group II intron, the reactive adenosine and two critical non-bridging phosphate oxygen atoms are separated by hundreds of nucleotides. The structural diversity surrounding these T-sites emphasizes that a wide range of tertiary structures will be detectable. T-site motifs consistently map to the center of important tertiary structure interactions or catalytic cores in our model RNAs.

Our whole-genome DENV T-site studies emphasize important features of T-site chemistry. First, T-site probing is highly selective. Only three strong T-sites were detected across the DENV RNA genome. Two T-sites display the specific TMO-reactive conformation in known 3D RNA structures^33,34^ and the third T-site occurs in a conserved structured region known to be functionally important^38^. Second, T-site chemistry identifies conserved tertiary structures despite sequence variation, as evidenced by our identification of T-sites in structurally conserved xrRNA and Dumbbell motifs, whose sequences vary between RNA virus classes. Third, the presence of a T-site reveals a tertiary structure with high specificity, but the absence of a T-site does not rule out a local tertiary structure. Fourth, RNA tertiary structures, specifically discoverable as T-sites, are harbingers of function as evidenced by the observation that all three DENV T-sites are functionally critical for viral replication. T-sites occur selectively at complex sites in RNA and such sites also comprise favored targets for small molecule ligands^40,41^. In sum, T-site chemistry is a facile, robust strategy that is poised to create broad opportunities to discover, analyze, and ligand RNA tertiary structures.

## Methods

### Electrostatics calculations

Solvated surface area and electrostatic potential for TMO and DMS were estimated using the MolCalc web server (molcalc.org)^42^.

### T-site detection thresholds

The reactivity threshold for T-site detection was determined empirically based on the TPP riboswitch, RNase P, and group II intron RNA data. Reactivities were measured as their direct fractional occurrence as measured by MaP. On this scale, the median reactivity for TMO and DMS across the three RNAs was 0.0085 and 0.0027and the median reactivity of the top 10% most reactive positions was 0.11 and 0.08, respectively. T-site signals were defined as the difference between TMO and DMS reactivities for each nucleotide. For this study, we required a T-site nucleotide to have a TMO over DMS signal of 0.2 or higher and for the ratio of TMO over DMS reactivity to be ≥3 (see Figure 2). This detection threshold was sufficient to discriminate against nucleotides that showed moderately enhanced TMO reactivity but did not show a T-site-like structure in known three-dimensional structures. For example, there is enhanced TMO reactivity at A63 in the RNase P RNA, but this position is base paired and does not adopt a T-site-like local structure. Varying (lowering) the T-site threshold does make it possible to identify additional T-sites, including in the DENV genomic RNA (Figure S6), with the expected cost of increased false-positive site identification.

### TMO handling

TMO was used as the tetrafluoroborate salt (MilliporeSigma). TMO stocks were handled in a fume hood, stored in a desiccator at −20 °C, and were used only once after opening. Stock solutions of TMO were prepared in 1:2 (vol/vol) nitromethane:sulfolane (from ThermoFisher and Sigma-Aldrich, respectively), as follows: TMO was first fully dissolved in nitromethane; subsequently, the TMO solution was diluted with two volumes sulfolane^21^. Final concentrations of TMO stock solutions used in this work (before addition of RNA) were 1.0 M. Sulfolane is a solid at room temperature and was warmed to 37 °C immediately prior to use. Due to the rapid kinetics of TMO hydrolysis (half-life ∼ 7 sec^21^), reactions were initiated by adding the larger volume of RNA solution to the smaller TMO volume, followed by mixing with immediate rapid pipetting.

### Synthesis of RNA constructs

DNA templates (IDT) for native and mutant sequences of the *Escherichia coli* TPP riboswitch (both aptamer domain and structure-switching constructs), the *Bacillus stearothermophilus* RNase P catalytic domain, and the *Oceanobacillus iheyensis* group II intron were embedded within 5’ and 3’ structure cassette flanking sequences and preceded with a 5’ T7 promoter sequence^43^. DNA templates were amplified by PCR [1 mL; 0.2 mM each dNTP, 500 nM each forward and reverse primer (IDT), 40 nM template, 0.02 U/µL DNA Polymerase (Q5 Hot Start High-Fidelity, NEB) using the temperature program: [95 °C for 30 s, 30 × (95 °C for 15 s, 65 °C for 30 s, 72 °C for 30 s), 72 °C for 2 min]. DNA template was recovered (Omega Mag-Bind beads, 1× ratio beads to PCR volume) and eluted in H_2_O. RNAs were transcribed in vitro [1 mL; 2.5 mM each NTP, 25 mM MgCl_2_, 40 mM Tris, pH 8.0, 2.5 mM spermidine, 0.01% (wt/vol) Triton X-100, 10 mM DTT, 0.025 units pyrophosphatase, 2.5 µg T7 polymerase, 1.125 µg DNA template, in 50 µL H_2_O; 37 °C; 6 h]. RNA was recovered (Omega Mag-Bind beads, 1.8× ratio beads to RNA volume) and resuspended in 50 µL of 0.1× TE (1 mM Tris pH 8.0, 0.1 mM EDTA).

### RNA folding and structure probing

Under the optimized conditions used here, both TMO and DMS react with all four ribonucleotides, with a preference for reaction with unpaired A and C residues^44^. The TPP riboswitch RNAs, RNase P, and group II intron RNA (5 pmol in 5 µL 0.1× TE for TPP constructs or in 6 µL 0.1× TE for RNase P and group II intron) were denatured at 95 °C for 2 min, cooled on ice, and incubated with 3 µL of 3× folding buffer (for TPP and RNase P, 33 mM MgCl_2_, 333 mM NaCl, 1 M bicine, pH 8; for the group II intron, 375 mM MgCl_2_, 333 mM NaCl, 1 M bicine, pH 8) at 37 °C for 20 min. If present, thiamine pyrophosphate ligand (1 µL of 50 µM thiamine pyrophosphate in H_2_O) was added to the TPP aptamer or structure-switching riboswitch samples, and all constructs were incubated at 37 °C for 10 additional minutes. After folding, the RNA solutions (9 µL) were added into 1 µL of reagent (100 mM TMO or 170 mM DMS final; both reagents in 1:2 v/v nitromethane-sulfolane). The TMO reaction is self-quenching; DMS was quenched with 1 µL of 2-mercaptoethanol after 6 min incubation at 37 °C^21^. The no-reagent control treatment contained 1 µL 1:2 v/v nitromethane-sulfolane. Modified RNA was purified (Illustra MicroSpin G-50 columns, GE Healthcare). No-Mg^2+^ structure probing was carried out exactly as described above but with MgCl_2_ omitted from the 3× folding buffer.

### MaP-RT

Modified RNAs (resuspended in TE) were subjected to reverse transcription essentially as described^39,44^. A 10 µL solution containing RNA, 200 nM gene specific primer, and 2 mM premixed dNTPs was incubated at 65 °C for 5 min followed by incubation at 4 °C for 2 min. To this solution was added 9 µL of 2.22× MaP buffer (1× MaP buffer contains 1 M betaine, 50 mM Tris, pH 8.0, 75 mM KCl, 10 mM DTT, 6 mM MnCl_2_), and the mixture was incubated at room temperature for 2 min. Reverse transcriptase (1 µL SuperScript II, Invitrogen) was added, and reverse transcription was performed using the following temperature program: 25 °C for 10 min, 42 °C for 90 min, 10 × [50 °C for 2 min, 42 °C for 2 min], 72 °C for 10 min. cDNA was then purified (Omega Mag Bind beads) and resuspended in H_2_O.

### Library preparation and sequencing

Sequencing libraries were prepared from cDNA products using a two-step PCR approach. In the first PCR step, 5-µL aliquots of purified cDNA were amplified for Illumina sequencing with the following temperature program: 98 °C for 30 s, 15 × [98 °C for 5 s, 68 °C for 20 s, 72 °C for 20 s], 72 °C for 2 min. The PCR products were recovered (Omega Mag-Bind beads) and eluted in water. In the second PCR step, treatment-specific barcodes were added to the ends of amplicons, with the following temperature program: 98 °C for 30 s, 10 × [98 °C for 5 s, 68 °C for 20 s, 72 °C for 20 s], 72 °C for 2 min. PCR products were recovered (Omega Mag-Bind beads), pooled, and sequenced. All libraries were sequenced on an Illumina MiSeq instrument using 300, 500, and 600 cycle kits for the TPP riboswitch, RNase P, and group II intron, respectively.

### Sequence alignment and mutation parsing

ShapeMapper (v2.1.5) was used to align reads to the reference sequences and to identify mutations created during MaP^45^. Default ShapeMapper settings were used to obtain profile.txt files, which are used downstream to analyze per-nucleotide reactivity data and to compare TMO and DMS reactivities. The --per-read-histograms option was used to generate high-level TMO and DMS reactivity distributions. The --output-parsed-mutations option was employed to produce mutation data for future ensemble deconvolution, if needed.

### DANCE deconvolution of TPP structure-switching RNA

For the TPP structure-switching riboswitch construct (only), TMO and DMS reactivities were normalized by dividing by the median reactivity of the top 10% most reactive positions. Ensembles for the structure-switching construct of the TPP riboswitch were deconvoluted with DanceMapper (v1.0)^31^. DANCE clustering was limited to a maximum of four components (-- maxc 4); in practice, DanceMapper identified two-state models as the best fit for experiments performed in the presence and absence of TPP. Structure cassette sequences were ignored and the first seven positions of the native riboswitch sequence were also ignored, to maintain sequence alignment with the simplified TPP riboswitch aptamer sequence^29^.

### Structure modeling

ShapeKnots (based on software package RNAstructure, v6.1) was used to model the secondary structures of the TPP aptamer, RNase P and group II intron RNAs^8,46^. Required parameters were the RNA primary sequences and the output.ct file name. Pseudo-free energy restraints were generated with the flag -dmsnt to use TMO- or DMS-normalized reactivities and the flag -x to incorporate PairMapper .bp files, thus introducing two TMO- or DMS-based bonuses (per-nucleotide and pairwise correlation) into the secondary structure calculation.^44^ The -m 1 option was used to select only the minimum free energy structure, outputting a .ct file containing the secondary structure model. foldClusters.py (part of DanceMapper) was used to model MFE structures (default options) for the structure-switching TPP riboswitch RNA.

### T-site and mutant reactivity analysis

Differential reactivity was assessed using reactivity profiles output by the ShapeMapper program. Per-nucleotide reactivity differences were computed by subtracting TMO reactivities from DMS reactivities. Ratios were computed similarly with TMO reactivities divided by DMS reactivities. Per-nucleotide reactivity profiles were visualized using RNavigate^47^. Three-dimensional structures of each RNA construct were visualized using PyMol (pymol.org). Individual motifs detected by differential probing were superimposed using the PyMol pair fitting function to identify conserved features. T-sites were defined as adenosine positions having ≥0.2 absolute reactivity difference between TMO and DMS, and a reactivity ratio of 2.5-fold for TMO versus DMS treatment, based on raw mutation rates. To identify the effect of T-site mutations, significant changes were defined as nucleotide positions having ≥0.05 absolute TMO reactivity difference between the mutant and native sequence RNA.

### DENV2 RNA structure probing

DENV2 (Dengue virus 2 New Guinea C; full name: Dengue virus 2 Thailand/NGS-C/1944) viral particles were purified from tissue culture supernatants from infected C6/36 cells as described^48^. Briefly, infected supernatant was layered over a 20% sucrose solution followed by ultracentrifugation. Virion pellets were then resuspended in 100 mM HEPES, pH 7.9 and 50 mM NaCl, followed by centrifugation. The DENV2 virion pellet was resuspended in 100 mM HEPES, pH 7.9 and RNA was extracted from purified virions (TRIzol LS reagent; ThermoFisher Scientific). The DENV2 RNA was then refolded in 10 mM MgCl_2_, 100 mM NaCl, 0.33 M Bicine, pH 8 and subjected to T-site chemical probing and MaP-RT as described above. cDNA from MaP-RT was subjected to second strand synthesis (non-directional second strand synthesis; NEBNext), followed by library preparation (Ultra II library prep kit; NEBNext). Two replicates, performed on separate days, were obtained and the same T-sites were observed in each experiment.

## Data Availability

Sequencing data are available from the Gene Expression Omnibus (GEO) at GSE286293.

## Supporting Information

One table of RNA sequences and six figures showing additional features of T-site chemistry.

## Acknowledgements

We are indebted to Amy Aponte Santana for insightful discussions. This work was supported by the US National Science Foundation (MCB-2027701 to K.M.W.), National Institutes of Health grants R01AI125416 (S.M.H) and T32GM142605 (C.J.A), and a National Science Foundation Graduate Research Fellowship (C.J.A).

