## Supporting Information for "A small cationic probe for accurate, punctate discovery of RNA tertiary structure"

[4] contributed equally

One table and six figures.

**Table S1. RNA sequences.** Structure cassette sequences are in lowercase.

| | |
|---|---|
| TPP Aptamer Native | ggccuucgggccaaGGACUCGGGGUGCCCUUCUGCGUGAAGGCUG<br>AGAAAUACCCGUAUCACCUGAUCUGGAUAAUGCCAGCGUAGG<br>GAAGUUCucgauccgguucgccggauccaaaucgggcuucgguccgguuc |
| TPP Aptamer U substitution | ggccuucgggccaaGGACUCGGGGUGCCCUUCUGCGUGAAGGCUG<br>AGAAAUACCCGUUUCACCUGAUCUGGAUAAUGCCAGCGUAGG<br>GAAGUUCucgauccgguucgccggauccaaaucgggcuucgguccgguuc |
| TPP Aptamer 3-nt deletion | ggccuucgggccaaGGACUCGGGGUGCCCUUCUGCGUGAAGGCUG<br>AGAAAUACCCGCACCUGAUCUGGAUAAUGCCAGCGUAGGGAA<br>GUUCucgauccgguucgccggauccaaaucgggcuucgguccgguuc |
| TPP Aptamer 1-nt deletion | ggccuucgggccaaGGACUCGGGGUGCCCUUCUGCGUGAAGGCUG<br>AGAAAUACCCGUUCACCUGAUCUGGAUAAUGCCAGCGUAGGG<br>AAGUUCucgauccgguucgccggauccaaaucgggcuucgguccgguuc |
| TPP structure switching construct | ggccuucgggccaaAACCAAACGACUCGGGGUGCCCUUCUGCGUGA<br>AGGCUGAGAAAUACCCGUAUCACCUGAUCUGGAUAAUGCCAGC<br>GUAGGGAAGUCACGGACCACCAGGUCAUUGCUUCUUCACGUU<br>AUGGCAGGAGCAAACUAUGCAAGUCGACCUGCUGGAUCCAGC<br>GCAAucgauccgguucgccggauccaaaucgggcuucgguccgguuc |
| RNase P Native | ggccuucgggccaaGUUAAUCAUGCUCGGGUAAUCGCUGCGGCCG<br>GUUUCGGCCGUAGAGGAAAGUCCAUGCUCGCACGGUGCUGAG<br>AUGCCCGUAGUGUUCGUGGAAACACGAGCGAGAAACCCAAAU<br>GAUGGUAGGGGCACCUUCCCGAAGGAAAUGAACGGAGGGAAG<br>GACAGGCGGCGCAUGCAGCCUGUAGAUAGAUGAUUACCGCCG<br>GAGUACGAGGCGCAAAGCCGCUUGCAGUACGAAGGUACAGAA<br>CAUGGCUUAUAGAGCAUGAUUAACGUCucgauccgguucgccggaucc<br>aaaucgggcuucgguccgguuc |
| RNase P U substitution | ggccuucgggccaaGUUAAUCAUGCUCGGGUAAUCGCUGCGGCCG<br>GUUUCGGCCGUAGUGGAAAGUCCAUGCUCGCACGGUGCUGAG<br>AUGCCCGUAGUGUUCGUGGAAACACGAGCGAGAAACCCAAAU<br>GAUGGUAGGGGCACCUUCCCGAAGGAAAUGAACGGAGGGAAG<br>GACAGGCGGCGCAUGCAGCCUGUAGAUAGAUGAUUACCGCCG<br>GAGUACGAGGCGCAAAGCCGCUUGCAGUACGAAGGUACAGAA<br>CAUGGCUUAUAGAGCAUGAUUAACGUCucgauccgguucgccggaucc<br>aaaucgggcuucgguccgguuc |
| RNase P 3-nt deletion | ggccuucgggccaaGUUAAUCAUGCUCGGGUAAUCGCUGCGGCCG<br>GUUUCGGCCGUAGAAAGUCCAUGCUCGCACGGUGCUGAGAUG<br>CCCGUAGUGUUCGUGGAAACACGAGCGAGAAACCCAAAUGAU<br>GGUAGGGGCACCUUCCCGAAGGAAAUGAACGGAGGGAAGGAC<br>AGGCGGCGCAUGCAGCCUGUAGAUAGAUGAUUACCGCCGGAG<br>UACGAGGCGCAAAGCCGCUUGCAGUACGAAGGUACAGAACAU<br>GGCUUAUAGAGCAUGAUUAACGUCucgauccgguucgccggauccaaau<br>cgggcuucgguccgguuc |

| | |
|---|---|
| RNase P 1-nt deletion | ggccuucgggccaaGUUAAUCAUGCUCGGGUAAUCGCUGCGGCCG GUUUCGGCCGUAGGGAAAGUCCAUGCUCGCACGGUGCUGAGA UGCCCGUAGUGUUCGUGGAAACACGAGCGAGAAACCCAAAUG AUGGUAGGGGCACCUUCCCGAAGGAAAUGAACGGAGGGAAGG ACAGGCGGCGCAUGCAGCCUGUAGAUAGAUGAUUACCGCCGG AGUACGAGGCGCAAAGCCGCUUGCAGUACGAAGGUACAGAAC AUGGCUUAUAGAGCAUGAUUAACGUCucgauccgguucgccggaucca aaucgggcuucgguccgguuc |
| RNase P loop mutant | ggccuucgggccaaGUUAAUCAUGCUCGGGUAAUCGCUGCGGCCG GUUUCGGCCGUAGAGGAAAGUCCAUGCUCGCACGGUGCUGAG AUGCCCGUAGUGUUCGUGGAAACACGAGCGAGAAACCCAAAU GAUGGUAGGGGCACCUUCCCGAAGGAAAUGUCCGGAGGGAAG GACAGGCGGCGCAUGCAGCCUGUAGAUAGAUGAUUACCGCCG GAGUACGAGGCGCAAAGCCGCUUGCAGUACGAAGGUACAGAA CAUGGCUUAUAGAGCAUGAUUAACGUCucgauccgguucgccggaucc aaaucgggcuucgguccgguuc |
| Group II Intron native | ggccuucgggccaaGUGUGCCCGGCAUGGGUGCAGUCUAUAGGGU GAGAGUCCCGAACUGUGAAGGCAGAAGUAACAGUUAGCCUAAC GCAAGGGUGUCCGUGGCGACAUGGAAUCUGAAGGAAGCGGAC GGCAAACCUUCGGUCUGAGGAACACGAACUUCAUAUGAGGCU AGGUAUCAAUGGAUGAGUUUGCAUAACAAAACAAAGUCCUUUC UGCCAAAGUUGGUACAGAGUAAAUGAAGCAGAUUGAUGAAGG GAAAGACUGCAUUCUUACCCGGGGAGGUCUGGAAACAGAAGU CAGCAGAAGUCAUAGUACCCUGUUCGCAGGGGAAGGACGGAA CAAGUAUGGCGUUCGCGCCUAAGCUUGAACCGCCGUAUACCG AACGGUACGUACGGUGGUGUGAGAGGAGUUCGCUCUACUCUA Uucgauccgguucgccggauccaaaucgggcuucgguccgguuc |
| Group II Intron U substitution | ggccuucgggccaaGUGUGCCCGGCAUGGGUGCAGUCUAUAGGGU GAGAGUCCCGAACUGUGAAGGCAGAAGUAACAGUUAGCCUAAC GCAAGGGUGUCCGUGGCGACAUGGAAUCUGAAGGAAGCGGAC GGCAAACCUUCGGUCUGAGGAACACGAACUUCAUAUGAGGCU AGGUAUCAAUGGAUGAGUUUGCAUAACAAAACAAAGUCCUUUC UGCCAAAGUUGGUACAGAGUAAAUGAAGCAGAUUGAUGAAGG GAAAGACUGCAUUCUUACCCGGGGAGGUCUGGAAACAGAAGU CAGCAGAAGUCAUAGUACCCUGUUCGCAGGGGAAGGACGGAA CAAGUAUGGCGUUCGCGCCUAAGCUUGAACCGCCGUAUACCG AACGGUUCGUACGGUGGUGUGAGAGGAGUUCGCUCUACUCUA Uucgauccgguucgccggauccaaaucgggcuucgguccgguuc |
| Group II intron 3-nt deletion | ggccuucgggccaaGUGUGCCCGGCAUGGGUGCAGUCUAUAGGGU GAGAGUCCCGAACUGUGAAGGCAGAAGUAACAGUUAGCCUAAC GCAAGGGUGUCCGUGGCGACAUGGAAUCUGAAGGAAGCGGAC GGCAAACCUUCGGUCUGAGGAACACGAACUUCAUAUGAGGCU AGGUAUCAAUGGAUGAGUUUGCAUAACAAAACAAAGUCCUUUC UGCCAAAGUUGGUACAGAGUAAAUGAAGCAGAUUGAUGAAGG GAAAGACUGCAUUCUUACCCGGGGAGGUCUGGAAACAGAAGU CAGCAGAAGUCAUAGUACCCUGUUCGCAGGGGAAGGACGGAA |

| | |
|---|---|
| | CAAGUAUGGCGUUCGCGCCUAAGCUUGAACCGCCGUAUACCG<br>AACGGGUACGGUGGUGUGAGAGGAGUUCGCUCUACUCUAUucg<br>auccgguucgccggauccaaaucgggcuucgguccgguuc |
| Group II intron 1-nt deletion | ggccuucgggccaaGUGUGCCCGGCAUGGGUGCAGUCUAUAGGGU<br>GAGAGUCCCGAACUGUGAAGGCAGAAGUAACAGUUAGCCUAAC<br>GCAAGGGUGUCCGUGGCGACAUGGAAUCUGAAGGAAGCGGAC<br>GGCAAACCUUCGGUCUGAGGAACACGAACUUCAUAUGAGGCU<br>AGGUAUCAAUGGAUGAGUUUGCAUAACAAAACAAAGUCCUUUC<br>UGCCAAAGUUGGUACAGAGUAAAUGAAGCAGAUUGAUGAAGG<br>GAAAGACUGCAUUCUUACCCGGGGAGGUCUGGAAACAGAAGU<br>CAGCAGAAGUCAUAGUACCCUGUUCGCAGGGGAAGGACGGAA<br>CAAGUAUGGCGUUCGCGCCUAAGCUUGAACCGCCGUAUACCG<br>AACGGUCGUACGGUGGUGUGAGAGGAGUUCGCUCUACUCUAU<br>ucgauccgguucgccggauccaaaucgggcuucgguccgguuc |
| Dengue virus 2 New Guinea C strain | DENV2 sequence used was Dengue virus serotype 2 New Guinea C. Complete sequence is available at https://www.ncbi.nlm.nih.gov/nuccore/KM204118.1 |

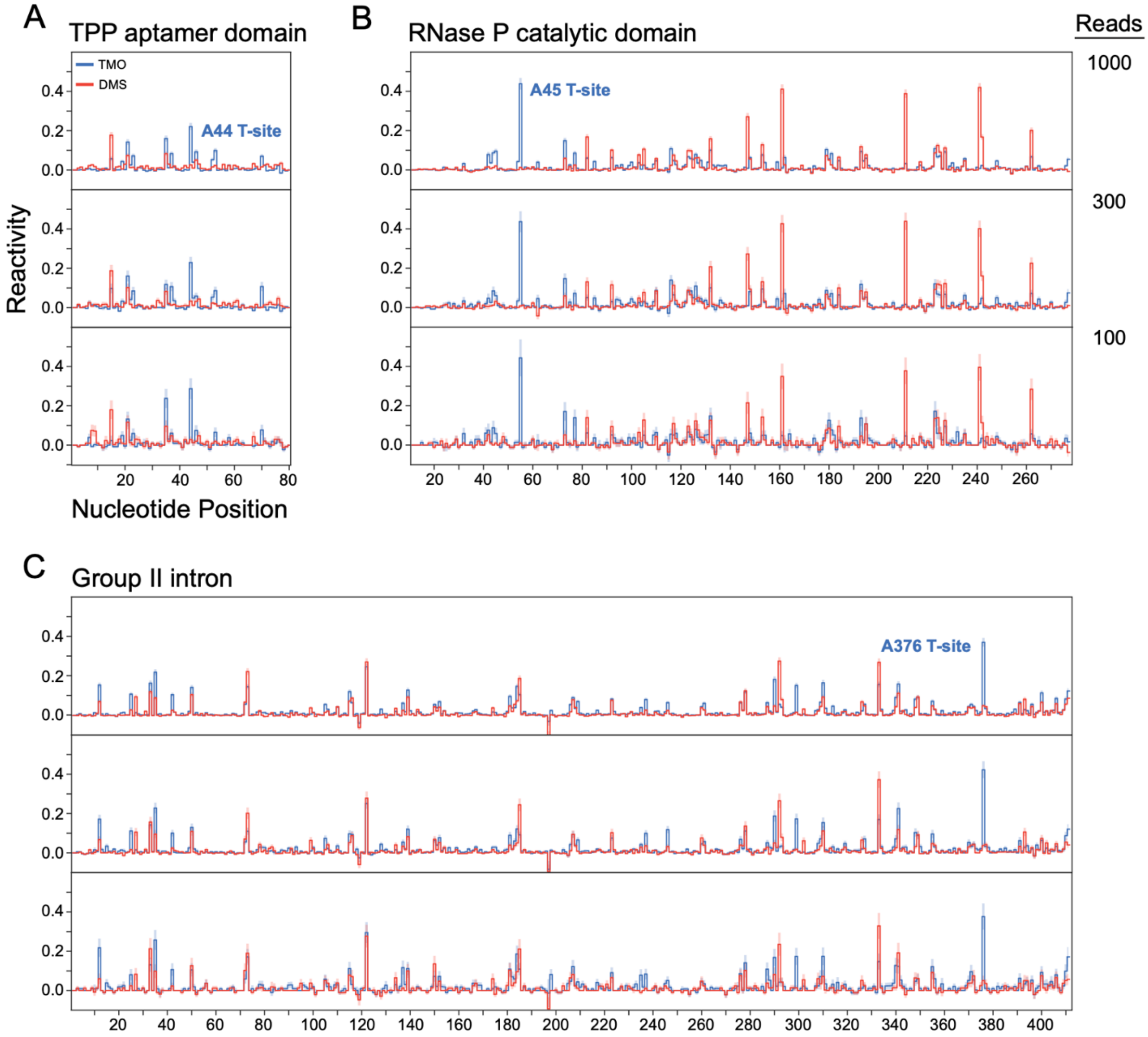


**Figure S1: Detection of T-sites at low sequencing read depths.** TMO (blue) and DMS (red) reactivity profiles for (**A**) TPP aptamer domain, (**B**) RNase P, and (**C**) group II intron RNAs at (down sampled) read depths of 1000 (top), 300 (middle), and 100 (bottom) reads. T-sites are labeled in bold text. Standard errors are shown as shaded bars for each nucleotide.

Note: T-site detection demonstrates high signal-to-noise, with the rate of TMO reactivity enhancement remaining consistently many-fold higher than the estimated error even at sequencing read depths of 100. T-site probing is thus appropriate for transcriptome-wide tertiary structure discovery.

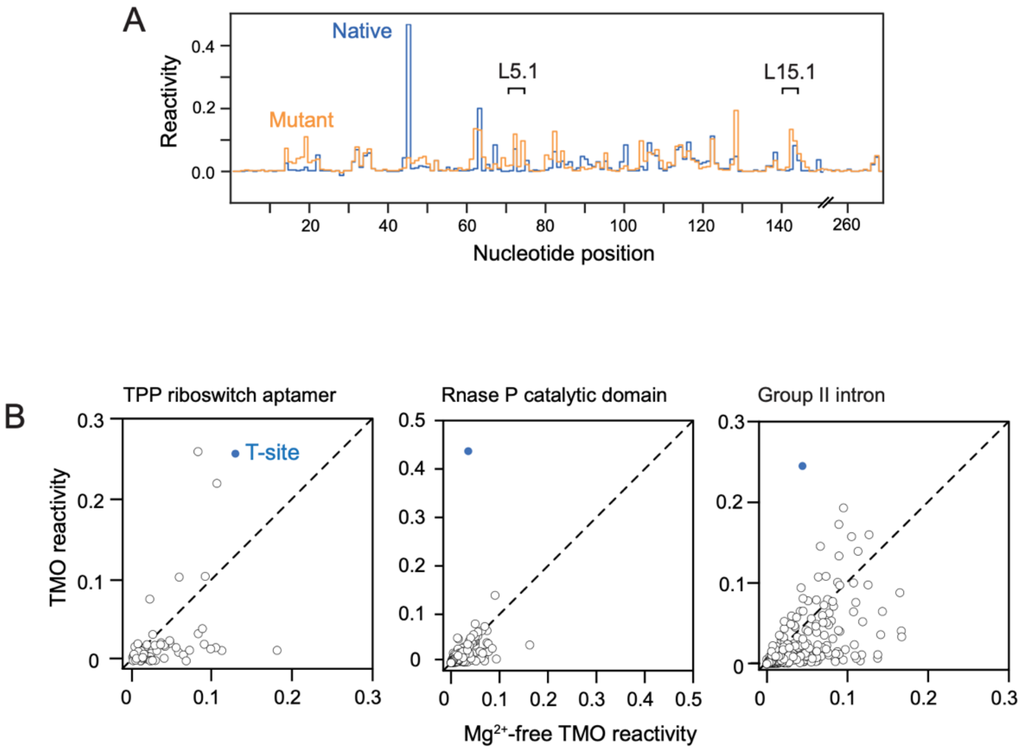


**Figure S2: Effect of tertiary structure disruption on T-site reactivity.** (**A**) Comparison of TMO reactivities for native sequence (blue) and tertiary structure disrupting mutant (orange). L5.1 and L15.1 regions are emphasized; the AA to CT tertiary-structure-disrupting mutation lies in L15.1. Critically, this mutant has the same complex secondary structure as the native RNase P RNA, but does not fold to create a tightly packed tertiary structure[21]. (**B**) TMO reactivities as a function of $Mg^{2+}$. $Mg^{2+}$ preferentially stabilizes RNA tertiary structure[14]. Raw TMO reactivities in the absence of $Mg^{2+}$ (x-axis) are plotted versus reactivities observed in standard $Mg^{2+}$-containing folding buffer. T-sites are shown with solid blue symbol.

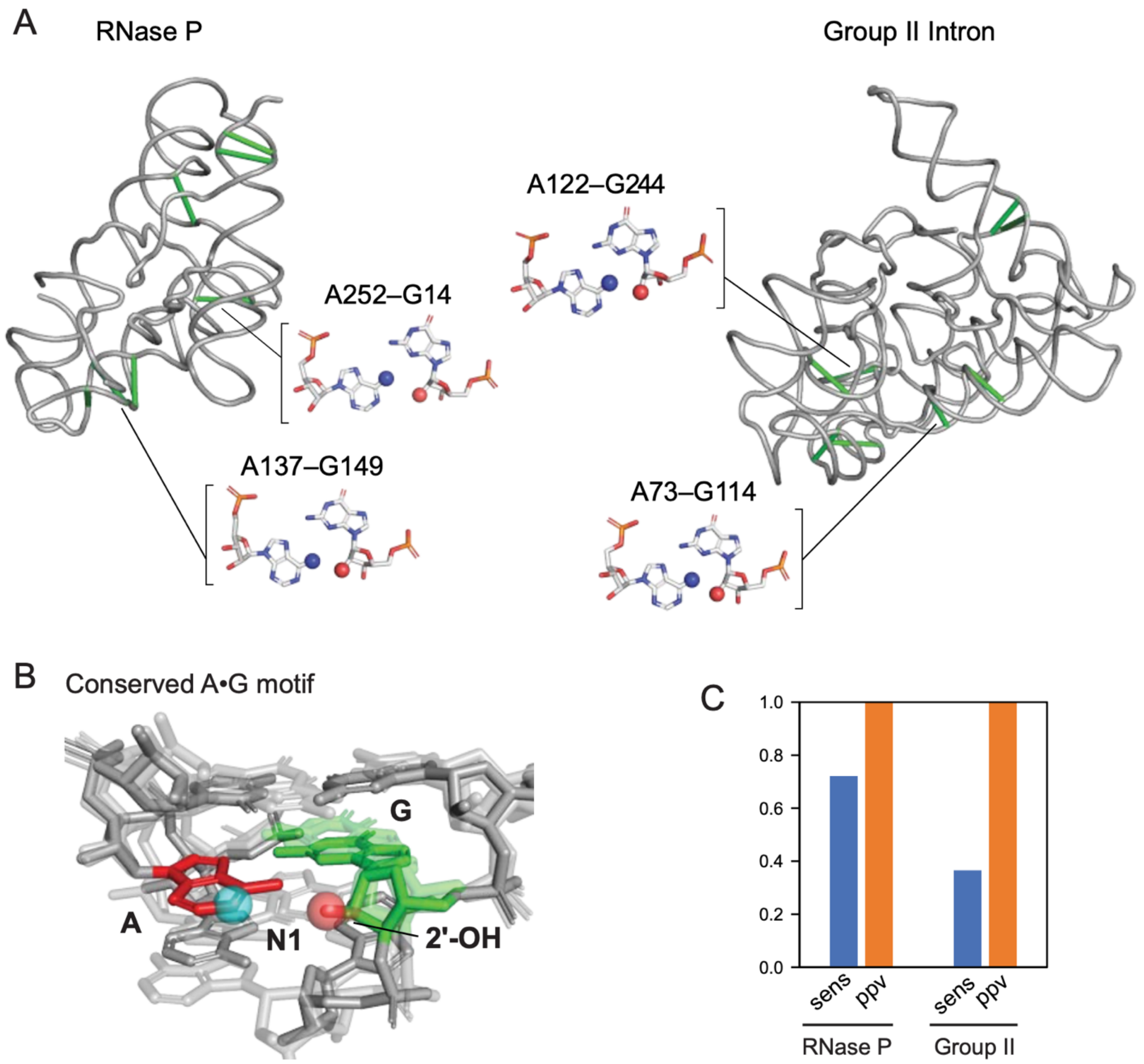


**Figure S3: Detection of A•G pairs as enhanced DMS reactivity.** (**A**) Ribbon diagrams of RNase P (left) and group II intron (right) structures. Green lines indicate Hoogsteen-sugar edge A•G pairs. Structures of representative pairs are illustrated. (**B**) Superimposition of four representative A•G base pairs reveal proximity of the N1 position in the reactive adenosine residue to the 2'-OH from the paired guanosine. (**C**) Sensitivity (sens) and positive predictive value (ppv) for detection of Hoogsteen-sugar edge A•G pairs using a DMS minus TMO difference rate threshold of 0.2.

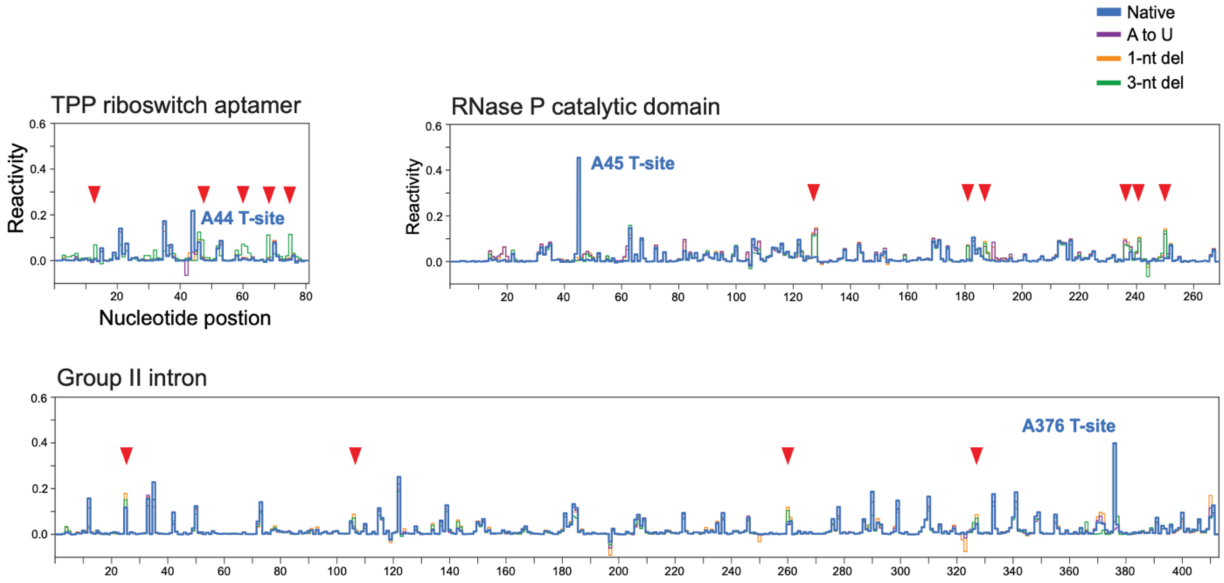


**Figure S4: Effect of mutating T-sites for TPP aptamer domain, RNase P, and group II intron RNAs.** TMO reactivity profiles for native and mutant sequences are shown. Nucleotides that show significant (≥0.05) reactivity increases in the 3-nt deletion mutants are denoted with red triangles. These data are visualized on three-dimensional structures in Figure 4.

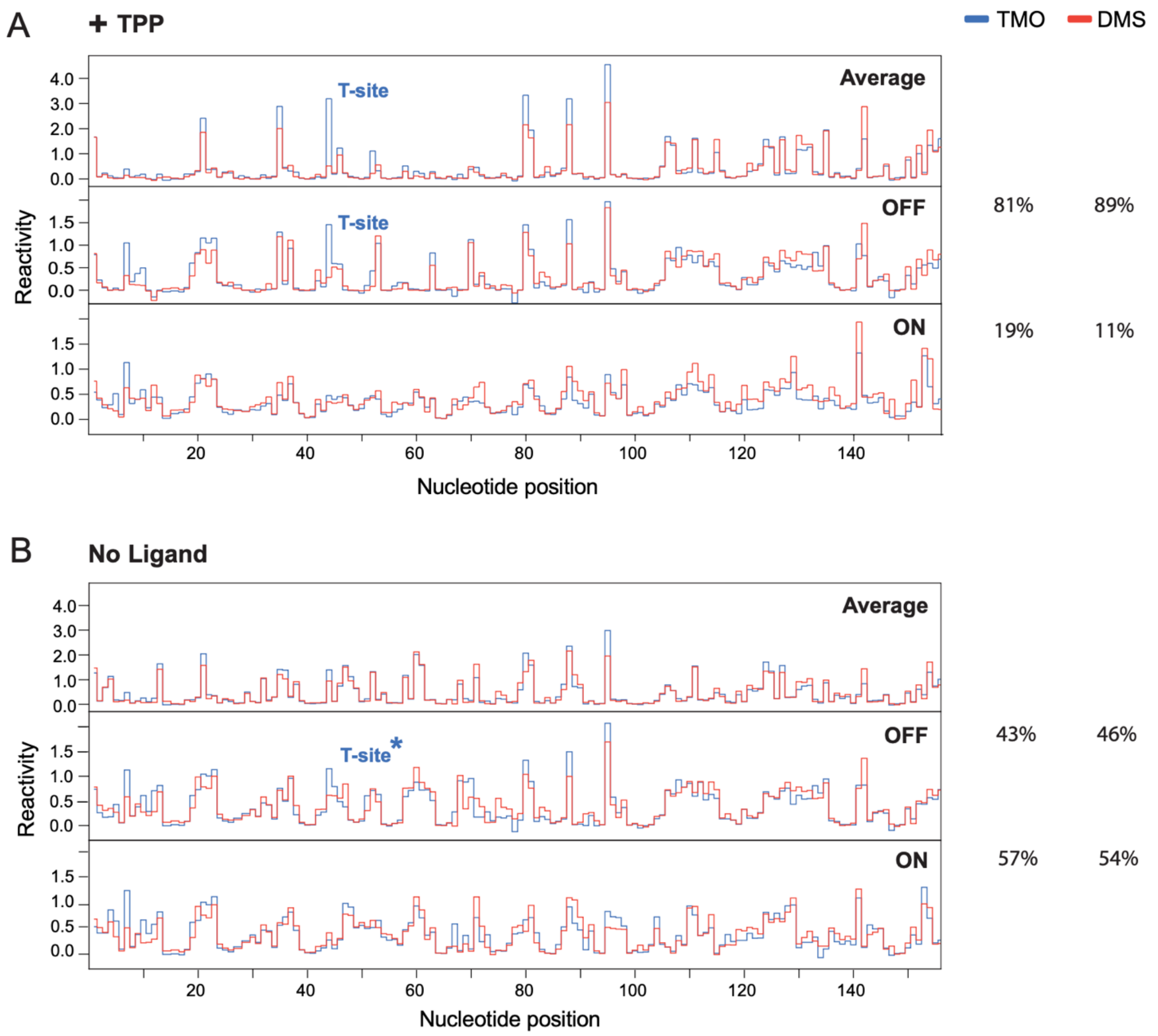


**Figure S5: Deconvolution of TMO and DMS reactivities in the presence and absence of ligand for the structure-switching TPP riboswitch.** TMO (blue) and DMS (red) reactivity profiles for the TPP riboswitch in the (**A**) presence or (**B**) absence of TPP ligand. Upper panels show averaged bulk reactivities, prior to deconvolution. Middle and lower panels show reactivities deconvoluted by DANCE-MaP. Population percentages for each translation state, as deconvoluted based on TMO or DMS reactivities, are listed at right. T-site at position A44, if present, is labeled in blue. Asterisk indicates T-site only visible in deconvoluted state. Data from panel A are reproduced from Figure 5.

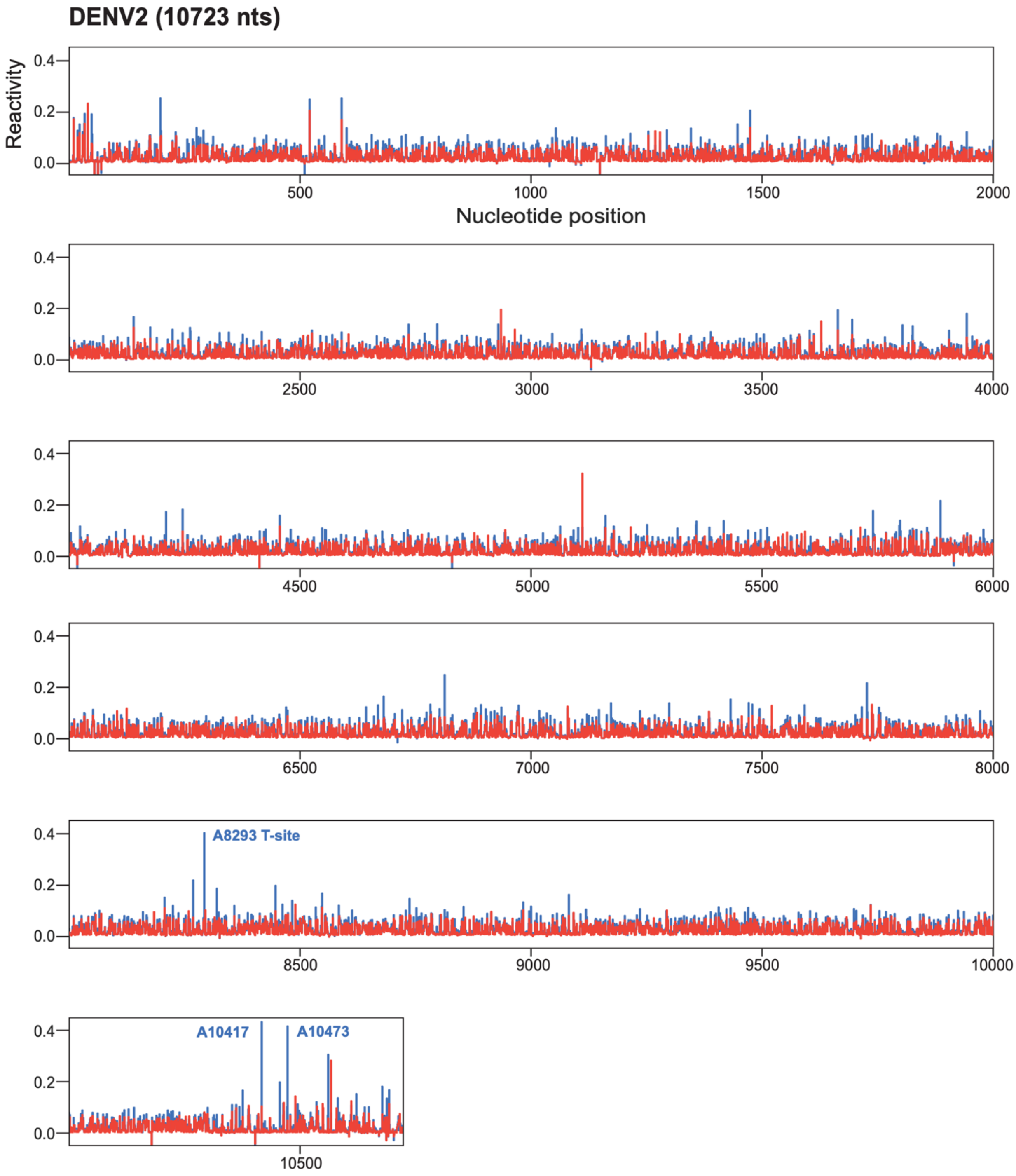


**Figure S6: T-site probing across the DENV2 RNA genome.** Reactivity profiles for TMO (blue) and DMS (red). Three mapped T-sites are labeled. The entire DENV2 RNA genome (New Guinea C strain; 10,723 nts) is shown; experiments were performed in replicate with one representative provided here.
